# Functional segregation of the human basal forebrain using resting state neuroimaging

**DOI:** 10.1101/211086

**Authors:** Ross D. Markello, R. Nathan Spreng, Wen-Ming Luh, Adam K. Anderson, Eve De Rosa

## Abstract

The basal forebrain (BF) is poised to play an important neuromodulatory role in brain regions important to cognition due to its broad projections and complex neurochemistry. While significant *in vivo* work has been done to elaborate BF function in nonhuman rodents and primates, comparatively limited work has examined the *in vivo* function of the human BF. In the current study we used multi-echo resting state functional magnetic resonance imaging (rs-fMRI) from 100 young adults (18-34 years) to assess the potential segregation of human BF nuclei as well as their associated projections. Bottom-up clustering of voxel-wise functional connectivity maps yielded adjacent functional clusters within the BF that closely aligned with the distinct, hypothesized nuclei important to cognition: the nucleus basalis of Meynert (NBM) and the medial septum/diagonal band of Broca (MS/DB). Examining their separate functional connections, the NBM and MS/DB revealed distinct projection patterns, suggesting a conservation of nuclei-specific functional connectivity with homologous regions known to be anatomically innervated by the BF. Specifically, the NBM demonstrated coupling with a widespread cortical network as well as the amygdala, whereas the MS/DB revealed coupling with a more circumscribed network, including the orbitofrontal cortex and hippocampal complex. Collectively, these *in vivo* rs-fMRI data demonstrate that the human BF nuclei support functional networks distinct aspects of resting-state functional networks, suggesting the human BF may be a neuromodulatory hub important for orchestrating network dynamics.

**Highlights:**

- The basal forebrain NBM and the MS/DB support two distinct functional networks
- Functional networks closely overlap with known anatomical basal forebrain
- Basal forebrain networks are distinct from known resting-state functional networks

## 1 Introduction

The basal forebrain (BF) serves as a major neuromodulatory hub for brain areas critical to cognition (Rye et al., 1984; Wenk, 1997; Yang et al., 2017). Due to its widespread projections (Descarries et al., 2004; Lysakowski et al., 1989; Mesulam et al., 1986), the BF has the capacity to play an important modulatory role in a range of neural dynamics (Baxter and Bucci, 2013; Hasselmo and Sarter, 2011; Ovsepian et al., 2016), influencing sensory plasticity (Picciotto et al., 2012; Rasmusson, 2000; Sarter and Bruno, 1997; Shinoe et al., 2005), attention (Botly and De Rosa, 2007, 2008, 2009, 2012; Gritton et al., 2016; Ljubojevic et al., 2014; Parikh et al., 2007), and memory (Chudasama et al. 2004; Croxson et al. 2011; cf. Baxter et al. 1995).

BF function has also been implicated in the cognitive changes associated with normal and pathological aging (Coyle et al., 1983; Davies and Maloney, 1976; Grothe et al., 2013; Muir, 1997; Perry et al., 1977; Whitehouse et al., 1982), and degeneration of the BF and its anatomical projections is hypothesized to have both functional and cognitive implications across the lifespan (Pepeu and Giovannini, 2017; Schmitz et al., 2016). A better understanding of human BF functional connections may provide insight into the distinct capacity of the BF to modulate the cortical and subcortical interactions that support cognition (Bell and Shine, 2016; Muñoz and Rudy, 2014).

Noninvasive assessments of human BF function have proven challenging due to difficulties in localization with functional magnetic resonance imaging (fMRI), especially when using whole-brain measurements. A stereotaxic probabilistic map of the BF generated using postmortem brains (Zaborszky et al., 2008) has partially ameliorated these issues (Li et al., 2014; Schmitz et al., 2016; Zhang et al., 2016). However, while significant work has demonstrated divisions in the neuro-modulatory projections of the BF in nonhuman animals with increasing specificity, to date no work has examined how the BF may functionally differentiate in humans (Golden et al., 2016; Hedreen et al., 1984; Kim et al., 2016, 2015; Kondo and Zaborszky, 2016; Ma and Luo, 2012; Mesulam et al., 1983b; Swanson and Cowan, 1979). It is thus important to assess the human functional networks mediated by the BF as it not only relates to existing knowledge of anatomical projections derived from these nonhuman animals investigations, but also to our current understanding of functional connections revealed through resting state fMRI data.

Rodent and nonhuman primate literature has shown the BF comprises distinct subdivisions, termed Ch1-4, delineating the BF at both a cellular and functional level (Mesulam et al., 1983b). The Ch1-3 groups, termed the medial septum and diagonal band of Broca (MS/DB), have projections that extend primarily into the hippocampal complex and hypothalamus (Hedreen et al., 1984; Swanson and Cowan, 1979), while the Ch4 group, termed the nucleus basalis of Meynert (NBM; also includes the substantia innominata), provides innervation to the entire cortical mantle and amygdala (Hedreen et al., 1984; Mesulam et al., 1983b). These subdivisons were originally differentiated based on their cholinergic projections (Mesulam et al., 1983b), but later work has demonstrated that the majority of projections from these regions are GABA-(Brashear et al., 1986; Gritti et al., 1993) and glutamatergic (Gritti et al., 2006). Indeed, co-release of GABA and glutamate from Ch1-4 projections is common (Gritti et al., 2006; Manns et al., 2001; Saunders et al., 2015), and recent work has shown that cholinergic and non-cholinergic projections from these BF functional subdivisions may work in tandem to influence cognition (Kim et al., 2015).

Thus, given the breadth and heterogeneity of its neuromodulatory connections, assessing how the BF relates to functional connectivity would be especially beneficial. Understanding how these neuromodulatory systems may influence functional connections and help inform whole-brain communication patterns is an especially relevant, yet understudied, aspect of the human fMRI literature (Bell and Shine, 2016; Shine et al., 2017; Spronk et al., 2017). The current study aimed to investigate the functional profiles of the BF using multi-echo resting-state fMRI (rsfMRI) data in young adults. Resting-state fMRI, in particular, has shown to be useful for delineating functional networks relevant to cognition (Stevens and Spreng, 2014), and recent developments in multi-echo fMRI have been shown to better separate BOLD responses tied to neural activity from non-BOLD artifacts (Kundu et al., 2013, 2017). We thus took advantage of a multi-echo approach to potentially distinguish activity within BF nuclei, along with their functional connectivity to known targets. We further compared traditional single-echo analyses with multi-echo techniques to assess the proportional dissociation of the MS/DB from the NBM. We hypothesized that rsfMRI would reveal functional networks reflecting known delineations of the BF, in support of cross-species homology previously demonstrated with *ex vivo* techniques (Emre et al., 1993; Mesulam et al., 1988; Saper and Chelimsky, 1984; Selden et al., 1998).

## 2 Materials and Methods

### 2.1 Participants

Participants were 100 young adults (18–34 years, mean = 22.6 ± 3.2 years, 58 females) with normal or corrected-to-normal vision and no history of psychiatric or neurological illness. All participants gave informed consent in accordance with the Institutional Review Board at Cornell University.

### 2.2 MRI data collection

Imaging data were acquired on a 3T GE Discovery MR750 MRI scanner with a 32-channel head coil at the Cornell Magnetic Resonance Imaging Facility in Ithaca, NY. Anatomical scans were captured using a T1-weighted volumetric MP-RAGE sequence (TR = 7.7 ms; TE = 3.42 ms; 7° flip angle; 1.0 mm isotropic voxels, 176 slices). Two 10 minute and 12 second BOLD resting state functional scans were acquired using a multi-echo EPI pulse sequence (TR = 3000 ms; TE_1_ = 13.7 ms, TE_2_ = 30.0 ms, TE_3_ = 47.0 ms; 83° flip angle; 3.0 mm isotropic voxels; 46 slices). Participants were instructed to remain awake with their eyes open for the duration of both functional scans.

### 2.3 MRI data preprocessing

All fMRI data were preprocessed in AFNI (version 16.0.12; Cox 1996), with the exception of skull-stripping and segmentation, which were done with FSL's Brain Extraction Tool (`bet2`; Jenkinson et al. 2005; Smith 2002) and FMRIB's Automated Segmentation Tool (`fast`; Zhang et al. 2001), respectively. Analysis was conducted using a combination of AFNI and Python (v3.5.3; https://www.python.org/) with associated NumPy (v1.11.1; Walt et al. 2011) and SciPy (v0.17.1; Jones et al. 2001) modules.

### 2.3.1 Multi-echo preprocessing

Multi-echo data were processed using ME-ICA (`meica.py`, version 2.5 beta 8; Kundu et al. 2012). The ME-ICA preprocessing pipeline was as follows: (1) slice timing correction was applied to all the TE series; (2) the first echo series (TE_1_) was aligned to the first time point using a 6-parameter rigid-body alignment procedure to generate motion correction parameters; (3) all TE series were used to generate a T2* map of the brain, which was co-registered to the skull-stripped anatomical to generate co-registration parameters; (4) the skull-stripped anatomical was linearly warped to MNI (Montreal Neurological Institute) space with the single-subject reference brain (Collins et al., 1994; Holmes et al., 1998) using a 12-parameter affine alignment procedure; (5) a brain mask was created from the mean image of the first echo (TE_1_) and applied to all the echoes; (6) motion correction, co-registration, and normalization warping parameters were simultaneously applied to each TE series; and (7) TE series were resampled to 2mm isotropic voxels.

The three TE series were then optimally combined to form a single, weighted time series. This combined time series was subjected to a FastICA decomposition and component sorting method (described in Kundu et al. 2013). Briefly, T2* signal in fMRI data linearly increases with TE, permitting classification of components as BOLD (T2*) or non-BOLD (S0) based on their adherence to a TE-dependent signal increase across the multiple echoes (see Kundu et al. 2017 for a recent review). Components classified as T2* BOLD were recombined to create a denoised time series. Multi-echo processed data were not smoothed further.

#### 2.3.2 Single-echo preprocessing

To compare the efficacy of ME-ICA in assessing functional connectivity, we separately preprocessed the middle echo image (i.e., TE_2_). The middle echo image was carried through the same processing pipeline detailed above through the end of step 7, at which point it was processed in a manner similar to Power et al. (2014).

Nuisance variables were generated by combining motion estimates (6), their derivatives (6), and tissue-specific estimates generated by averaging signal across voxels in tissue-specific masks (produced by FSL's fast). In all, six tissue-specific estimates were used: cerebrospinal fluid (CSF), white matter (WM), and the global signal (whole brain), as well as their derivatives.

Frame-wise displacement (FD) and change in signal variance (DVARS, **D**erivative of signal **VAR**iance) traces were calculated from the middle echo time series using tools implemented in Nipype (Gorgolewski et al., 2011); visual inspection of these traces across subjects suggested that FD ≥ 0.2mm and DVARS ≥ 1.6% ΔBOLD were adequate cutoffs for identifying abnormal motion. Following Power et al. (2012), temporal masks were created from each of these traces: frames that exceeded the threshold were marked for removal, along with the frame before and the two frames after the marked timepoint. The intersection of the temporal masks generated from each trace were used to create a final mask.

The final temporal mask was applied to all nuisance variables and the TE_2_ time series; standardized and detrended nuisance variables were regressed against the time series, generating least-squares fit beta maps. Beta maps were multiplied with the “full” (i.e., unmasked) nuisance variables to generate a modeled time-series, which was subtracted from the observed time series. The resultant residual time series was quadratically detrended, bandpass filtered (0.009-0.08 Hz) and smoothed (6mm FWHM) inside a whole brain mask (generated in step 5 of multi-echo preprocessing, above). Finally, the temporal mask was reapplied to the smoothed data; the first and last 13 TRs of each run were also removed to account for temporal smoothing due to filtering.

### 2.4 Regions of interest

Regions of interest (ROIs) were generated from the BF probabilistic maps in the SPM Anatomy toolbox (Eickhoff et al., 2007; Zaborszky et al., 2008). A whole BF mask was created by combining the probability masks, resampling to functional space, and thresholding at >40% probability. Individual ROI masks were created via the same procedure using the separate probability masks. Resultant masks were 95 voxels in size for entire BF (Ch1-4, BF), 37 voxels for MS/DB (Ch1-3), and 51 voxels for NBM (Ch4).

#### 2.4.1 Basal forebrain T2* values

The T2* value of each BF nucleus was calculated using the map generated during step (3) of the multi-echo data processing pipeline. Values were averaged within each ROI prior to being averaged across the two resting states runs for each individual. Resultant values were averaged across individuals to generate optimal T2* values for the NBM (29.97ms ± 4.48) and the MS/DB (28.63ms ± 9.94). The standard deviation of T2* values were also calculated for within the NBM (9.02ms ± 1.71) and within the MS/DB (10.63ms ± 2.62), suggesting some inhomogeneity within ROIs. In general, T2* values for both regions seem to align with echo times traditionally used in resting-state functional analyses.

#### 2.4.2 Quality assessment

Previous work has demonstrated the efficacy of ME-ICA in removing physiological and motion-related noise (Dipasquale et al., 2017; Kundu et al., 2015) and increasing temporal SNR (DuPre et al., 2016). However, given the proximity of the BF to regions which typically experience signal dropout and distortion, we wanted to assess the relative tSNR of each ROI to ensure adequate ability to detect effects of interest. Using AFNI's `3dTstat -cvarinvNOD`, tSNR was calculated for both multi-and single-echo preprocessed data inside each ROI for individual subjects according to:

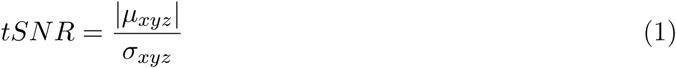

where *µ*_*xyz*_ is the average signal within a given voxel and *σ*_*xyz*_ is the standard deviation.

Subject-level tSNR values were averaged to generate an approximate group-level tSNR metric (Figure 1) for the NBM (multi-echo: 254.97 ± 129.01, single-echo: 68.01 ± 49.43) and MS/DB (multi-echo: 380.48 ± 149.44, single-echo: 111.28 ± 46.56). Given the use of global signal regression on single-echo data during preprocessing, tSNR for single-echo data was calculated on a more minimally preprocessed TE_2_ time series following conventional denoising performed in Kundu et al. (2013). Calculation of tSNR for multi-echo data was performed using the time series generated by projecting the BOLD-selected components into the time domain (referred to as the high-*κ* time series, Kundu et al. 2013).

**Figure 1.**
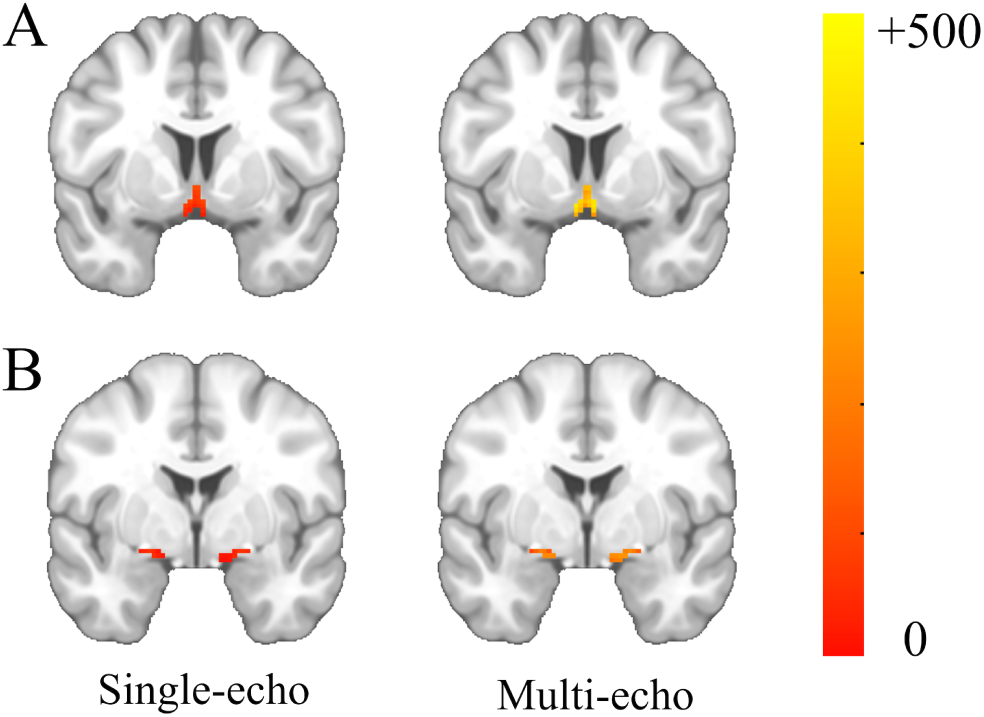
Temporal signal-to-noise ratio. **A**. The medial septum/diagonal band of Broca, and **B**. the nucleus basalis of Meynert for both single- and multi-echo data. Temporal signal-to-noise was calculated as the absolute value of the mean divided by the standard deviation of the time series, as in Equation 1. Figure made with FSLeyes software.

### 2.5 Functional connectivity

Multi-echo independent components regression (ME-ICR; Kundu et al. 2013) was used to assess functional correlations for multi-echo data; ME-ICR has been demonstrated to reliably capture known connections at the single-subject level with single-voxel coordinates (Kundu et al., 2013). Traditional voxel-wise, time-series correlations were used to assess functional relationships for single-echo data.

### 2.5.1 Multi-echo rsFC

Rather than using the BOLD time series, weights for voxels in each ROI were extracted from the independent BOLD components identified during the ME-ICA component sorting (described above) to ensure correlated activity represented BOLD-specific functional coupling. The Pearson correlation was calculated between the extracted vector of weights and that of every other voxel in the brain. Correlation maps were converted to Z scores using Fisher's *r*-to-*Z* transform:

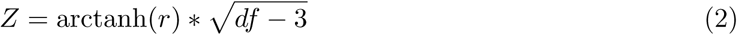

where *df* is the degrees of freedom—in this case, the number of BOLD components (i.e., the length of the weight vector), which varied across individuals and sessions. Z-score maps were averaged across both functional scans for a given subject before being brought to a group-level analysis.

We conducted separate, one-sample t-tests for the whole BF, NBM, and MS/DB Z-score maps. Group analyses were confined to a gray matter mask derived using FSL's fast on the single-subject MNI reference template resampled to functional space, and group results were smoothed with a 6mm FWHM Gaussian kernel.

### 2.5.2 Single-echo rsFC

Signal for voxels in each ROI were extracted and averaged, then correlated against every voxel in the brain. Correlation maps were converted to Z-score using Equation 2. Degrees of freedom were calculated with:

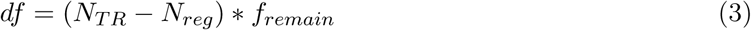

where *N*_*TR*_ is the number of TRs that remained after the final temporal mask was applied, *N*_*reg*_ is the number of nuisance regressors (here, 18), and *f*_*remain*_ is the proportion of frequencies remaining after bandpass filtering (here, 0.426), as in Kundu et al. (2013) and Power et al. (2014). Group level analyses were performed on resultant Z-score maps as described for multi-echo data.

Only positive findings are presented for single-echo results due to the potential introduction of artifactual anti-correlations through the use of global signal regressions (Murphy et al., 2009; Saad et al., 2012; Van Dijk et al., 2010; Weissenbacher et al., 2009). ME-ICR has been shown to avoid the skewed distribution of correlation coefficients found in traditional connectivity analyses (Kundu et al., 2013), thus supplanting the need for additional corrective measures with multi-echo data.

### 2.6 Ward clustering

To examine whether the functional connectivity profiles were segregated by BF nuclei, we employed hierarchical clustering using Euclidean distance and Ward's criterion (implemented with scipy; Jones et al. 2001) on voxel-wise rsFC values computed from the whole BF ROI mask (95 voxels). Ward clustering was selected over other clustering mechanisms due to previous reports of its superior accuracy and reproducibility in parcellating functional MRI data (Thirion et al., 2014). We were particularly interested in a *k* =2 cluster solution to directly examine if the two anatomical nuclei were the predominant, discriminable clusters within the functional data; however, solutions were generated for all cluster sizes. The most parsimonious solution was selected by choosing the one with the maximum silhouette score (implemented with scikit-learn; Pedregosa et al. 2011). Silhouette scores provide a metric for how similar elements of a given cluster are to each other compared to elements from other clusters; a higher silhouette score suggests a more appropriate clustering solution (Rousseeuw, 1987). Clusters were independently generated for multi-and single-echo data.

### 2.7 Partial correlations

To ensure that contributions of each BF nucleus to their derived functional connectivity profiles were unique, we conducted partial correlations on each nucleus, linearly regressing out signal from the other. To examine shared connectivity between the nuclei, we used a simple conjunction (boolean “AND”) on the results of these analyses (Nichols et al., 2005). Distinct connectivity patterns for each nucleus were examined with a simple disjunction (boolean “NOT”) of the shared patterns.

### 2.8 Thresholding

Group-level functional connectivity maps are presented as Z-scores, thresholded at *p* ≤ 0.001 uncorrected and cluster-corrected to *p* ≤ 0.01. Given the sample size of the current study, using an uncorrected *p* ≤ 0.001 threshold ensured all results were also significant at the FDR *q* ≤ 0.05 threshold. Group analyses and cluster correction for functional connectivity maps were implemented using AFNI's `3dttest++` command with the `-clustim` option, which generates approximately 10,000 permuted versions of the group-level residuals to calculate appropriate cluster thresholds based on intrinsic smoothness of the data. Despite recent concerns with this implementation of Type I error correction (Eklund et al., 2016), this method of cluster-correction appropriately attenuates the family-wise error rate to under 0.05 (Cox et al., 2016). All group-level results are presented using FSLeyes software (https://git.fmrib.ox.ac.uk/fsl/fsleyes/fsleyes). Inflated statistical maps can be found in Figure 5; full statistical maps can be viewed online at NeuroVault (https://neurovault.org/collections/QAMPLTIB).

## 3 Results

### 3.1 Basal forebrain resting-state functional connectivity

We assessed the functional connectivity profiles of the BF in healthy adults using multi-echo independent components regression and traditional single-echo time-series functional correlations. Group-level effects were calculated with independent one-sample t-tests.

Functional connectivity of the BF was first examined with ME-ICR, which computes the correlation between BOLD-identified component weight vectors generated via ME-ICA. Results revealed broad relationships with all of insulo-opercular cortex, the cingulum band, medial prefrontal cortex, and bilateral fusiform gyrus (see Figure 2A). Additionally, strong connections were present between the NBM and subcortical targets, including the amygdala and thalamus, extending into the cerebellum. A full list of regions can be found in Table 2.

**Figure 2.**
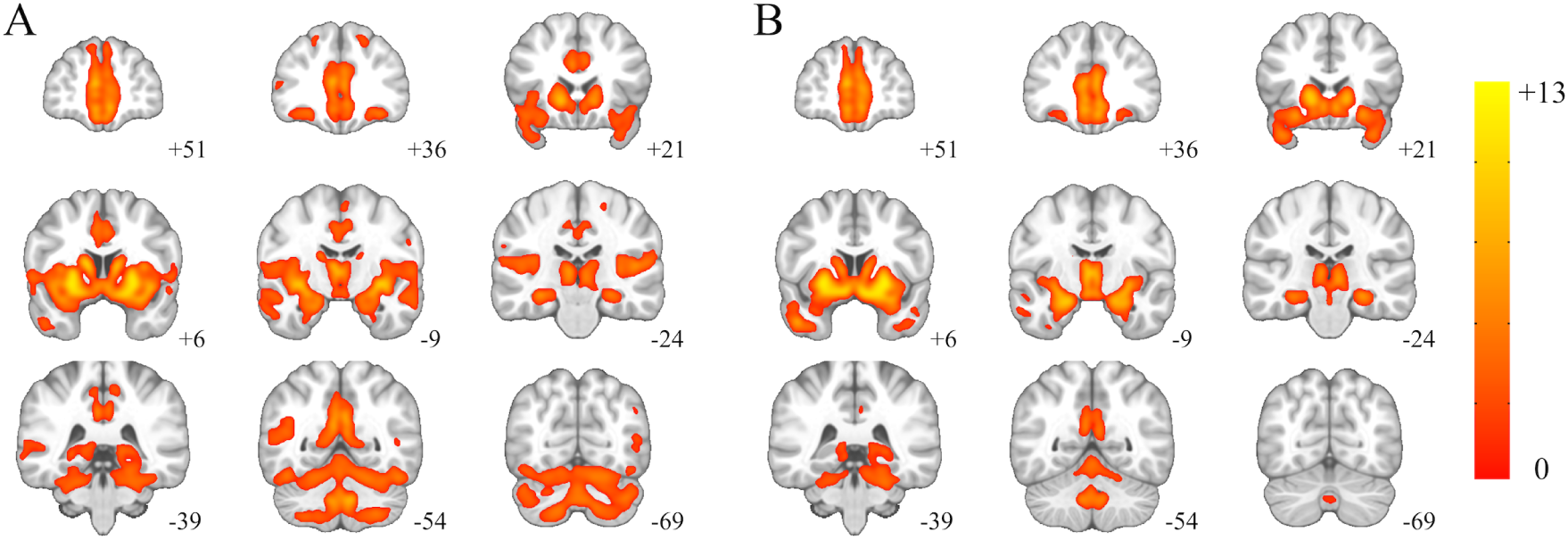
Functional correlations of the basal forebrain (BF, Ch1-4). **A**. Multi-echo preprocessed and **B**. single-echo preprocessed data. Significant connections are presented at a p ≤ 0.001threshold, cluster-corrected to alpha ≤ 0.01, and presented as Z-scores. Results are presented on the MNI-152 reference image to aid in anatomical localization.

Resting state functional connectivity (rsFC) of the BF was also assessed using traditional time-series correlations with single-echo processed data (Figure 2B). Similar connections were found between the BF and prefrontal cortex, insulo-opercular cortex, and subcortical features (i.e., the amygdala; Table 2).

### 3.2 Functional clustering

We used Ward clustering to examine the segregation of the BF nuclei based on functional connectivity and the potential correspondence with anatomy. Voxel-wise rsFC maps for both single-and multi-echo data were calculated for every voxel in the whole BF ROI against the rest of the brain and subjected to Ward clustering. Notably, for both single (0.3424) and multi-echo data (0.2275) silhouette scores were highest for a *k* =2 solution; these clustering solutions were carried forward for further analysis (Figure 3). Overlap with anatomical nuclei was calculated using the Sorensen-Dice index (Zou et al., 2004).

**Figure 3.**
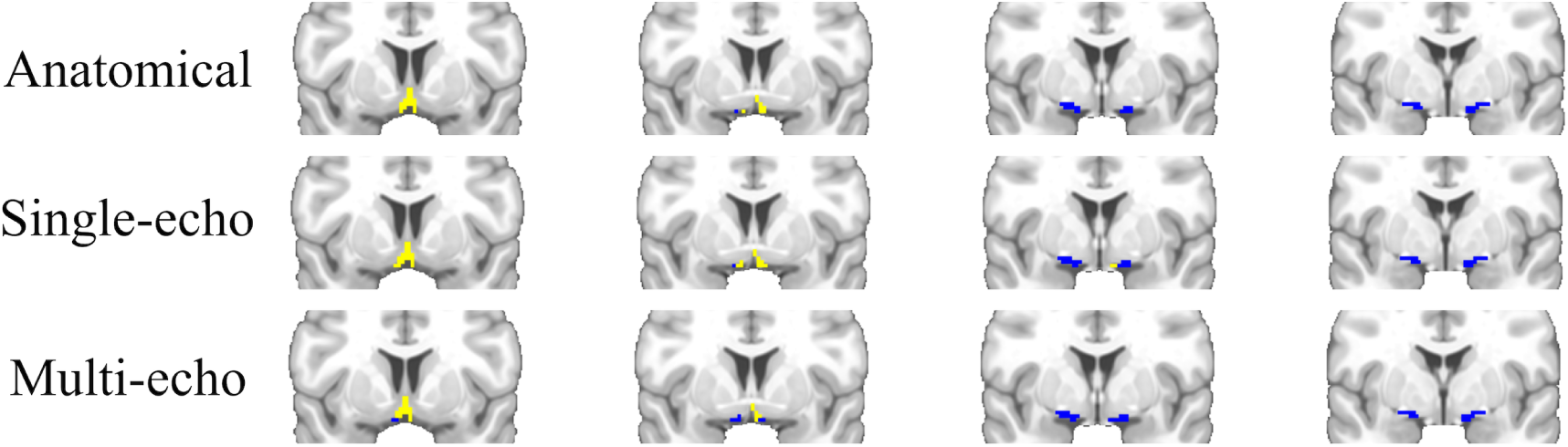
Anatomical and functional clusters of the basal forebrain. **Top**. Anatomical BF region of interest generated from Zaborszky et al. (2008) probability maps. **Center**. Cluster solution from ME-ICR functional connectivity data. **Bottom**. Cluster solution from single-echo functional connectivity data. Yellow corresponds to c1, blue to c2 in Table 1.

Multi-echo data generated two clusters of size 34 (c1) and 61 (c2) voxels, with very high affinity to anatomical delineations of BF nuclei (see Table 1 for Dice coefficients). Single-echo data resulted in clusters of size 43 (c1) and 52 (c2) voxels, respectively (Table 1).

**Table 1.**
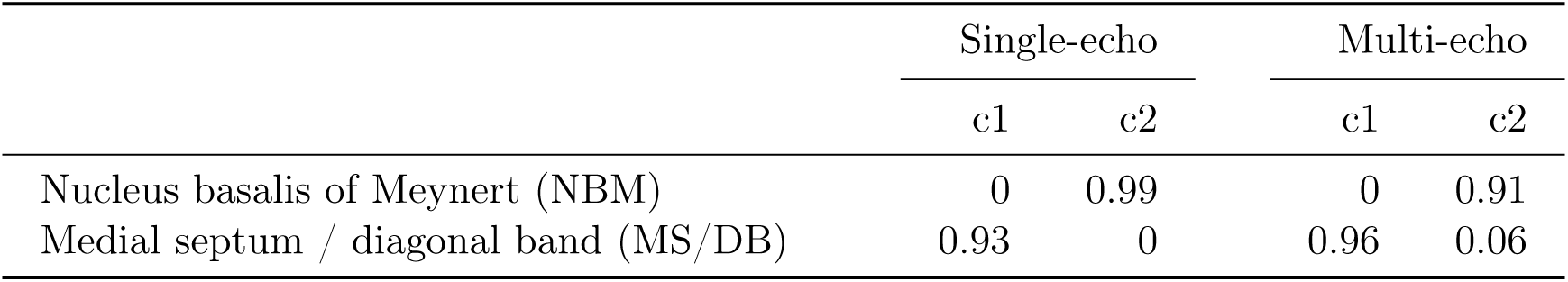
Dice coefficients of functional clusters and BF nuclei. Functional clusters were generated via Ward clustering using voxel-wise resting-state functional connectivity maps from the BF with a *k* =2 cluster solution. Functional cluster overlap with BF nuclei was calculated using the Dice coefficient. Labels c1 and c2 correspond respectively to yellow and blue clusters in Figure 3.

|  | Single-echo |  | Multi-echo |  |
| --- | --- | --- | --- | --- |
|  | c1 | c2 | c1 | c2 |
| Nucleus basalis of Meynert (NBM) | 0 | 0.99 | 0 | 0.91 |
| Medial septum / diagonal band (MS/DB) | 0.93 | 0 | 0.96 | 0.06 |

**Table 2.** Basal forebrain functional connectivity. Regions functionally connected with the BF as assessed via ME-ICR (multi-echo) and time-series rsFC (single-echo). Coordinates are provided in MNI (Montreal Neurological Institute) space. *k* values represent the voxel extent of the given cluster and *Z* values are from the peak coordinate in the cluster. Sub-peaks were assessed within larger clusters based on a |*Z*| > 10 threshold and a minimum distance ≥ 10mm from other sub-peaks; locations for sub-peaks are italicized.

| | $x$ | $y$ | $z$ | $k$ | $Z$ |
| --- | --- | --- | --- | --- | --- |
| *Multi-echo results* |  |  |  |  |  |
| Right ventral striatum | 23 | 10 | -5 | 45800 | 12.65 |
| *Left putamen* | -23 | 10 | -3 |  | 12.42 |
| *Right caudate nucleus* | 13 | 14 | 7 |  | 11.41 |
| *Right caudate nucleus* | 11 | 16 | -5 |  | 11.33 |
| *Left caudate nucleus* | -7 | 14 | -7 |  | 10.98 |
| *Right putamen* | 33 | -4 | -5 |  | 10.96 |
| *Left putamen* | -33 | -6 | -5 |  | 10.78 |
| *Left amygdala* | -23 | 0 | -15 |  | 10.17 |
| *Left caudate nucleus* | -11 | 10 | 7 |  | 10.13 |
| Left angular gyrus | -45 | -62 | 29 | 412 | 4.65 |
| Left superior frontal gyrus | -19 | 38 | 45 | 113 | 4.67 |
| Left lingual gyrus | -3 | -86 | -7 | 70 | 4.17 |
| Right precentral gyrus | 25 | -28 | 61 | 58 | 4.29 |
| Left inferior frontal gyrus | -37 | 28 | -1 | 56 | 4.14 |
| Right middle frontal gyrus | 39 | 18 | 25 | 54 | 4.25 |
| Left precentral gyrus | -21 | -26 | 59 | 37 | 4.20 |
| *Single-echo results* |  |  |  |  |  |
| Right ventral striatum | 23 | 10 | -5 | 23106 | 12.74 |
| *Left putamen* | -23 | 8 | -5 |  | 12.63 |
| *Right caudate nucleus* | 11 | 12 | -11 |  | 12.23 |
| *Left caudate nucleus* | -9 | 10 | -11 |  | 12.08 |
| *Right amygdala* | 23 | -4 | -17 |  | 12.00 |
| *Left caudate nucleus* | -9 | 16 | -5 |  | 11.91 |
| *Right caudate nucleus* | 13 | 18 | -1 |  | 11.77 |
| *Left amygdala* | -21 | -4 | -17 |  | 11.74 |
| *Right thalamus* | 1 | -12 | 5 |  | 11.13 |
| *Left hippocampus* | -27 | -14 | -19 |  | 10.90 |
| *Left anterior cingulate cortex* | -5 | 46 | 7 |  | 10.46 |
| *Left anterior cingulate cortex* | -5 | 46 | -1 |  | 10.40 |
| *Hypothalamus* | 7 | 2 | -11 |  | 10.26 |

An additional clustering analysis was done focusing solely on intra-BF connectivity: rsFC maps from within the BF were used as the input to a *k* =2 clustering. Results were similar to the whole-brain analysis: multi-echo data yielded two clusters of size 49 (NBM = 0.94, MS/DB = 0) and 46 (NBM = 0.08, MS/DB = 0.89) voxels respectively, while single-echo yielded clusters of size 46 (NBM = 0.93, MS/DB = 0) and 49 (NBM = 0.12, MS/DB = 0.86) voxels. The clusters followed anatomical delineations moderately well, but less so than those generated via whole-brain voxel-wise clustering.

### 3.3 Basal forebrain nuclei rsFC

Given the alignment of functional clustering of the BF with anatomical divisions, we independently examined potentially distinct functional profiles of the BF nuclei: the NBM and the MS/DB. As the functional parcels derived via Ward clustering demonstrated high correspondence with the anatomical ROIs (see Table 1), we opted to use the latter in order to examine the individual profiles of the subnuclei. Importantly, using the anatomical ROIs allowed us to continue to investigate the similarities between the single-and multi-echo analyses without biasing results.

In order to better assess shared and distinct functional correlations between the NBM and MS/DB, partial correlations were conducted on each nucleus, regressing out signal from the other, to ensure contributions were unique to the nucleus of interest.

#### 3.3.1 Shared NBM and MS/DB connectivity

A simple conjunction between the NBM and MS/DB functional correlation maps revealed regions with shared connectivity between the two BF nuclei (Figure 4; green regions). In multi-echo data, these connections are primarily found in the hippocampus, posterior cingulate cortex, and ventromedial prefrontal cortex (vmPFC), whereas overlap is localized to the hippocampus and caudate in single-echo data.

**Figure 4.**
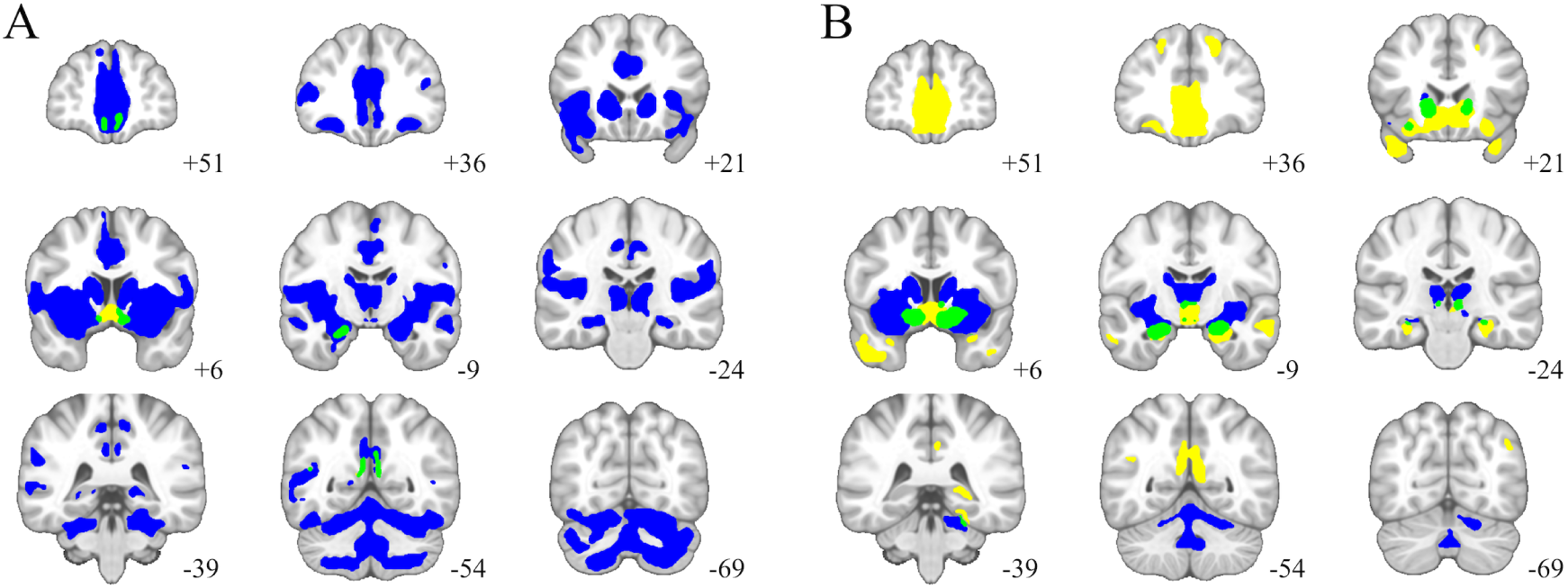
Functional correlations of the medial septum and diagonal band (MS/DB, Ch123) and nucleus basalis of Meynert (NBM, Ch4) for **A**. multi-echo preprocessed and **B**. single-echo preprocessed data. Yellow denotes connectivity specific to the MS/DB, blue specific to the NBM, and green is shared connectivity.

**Figure 5.**
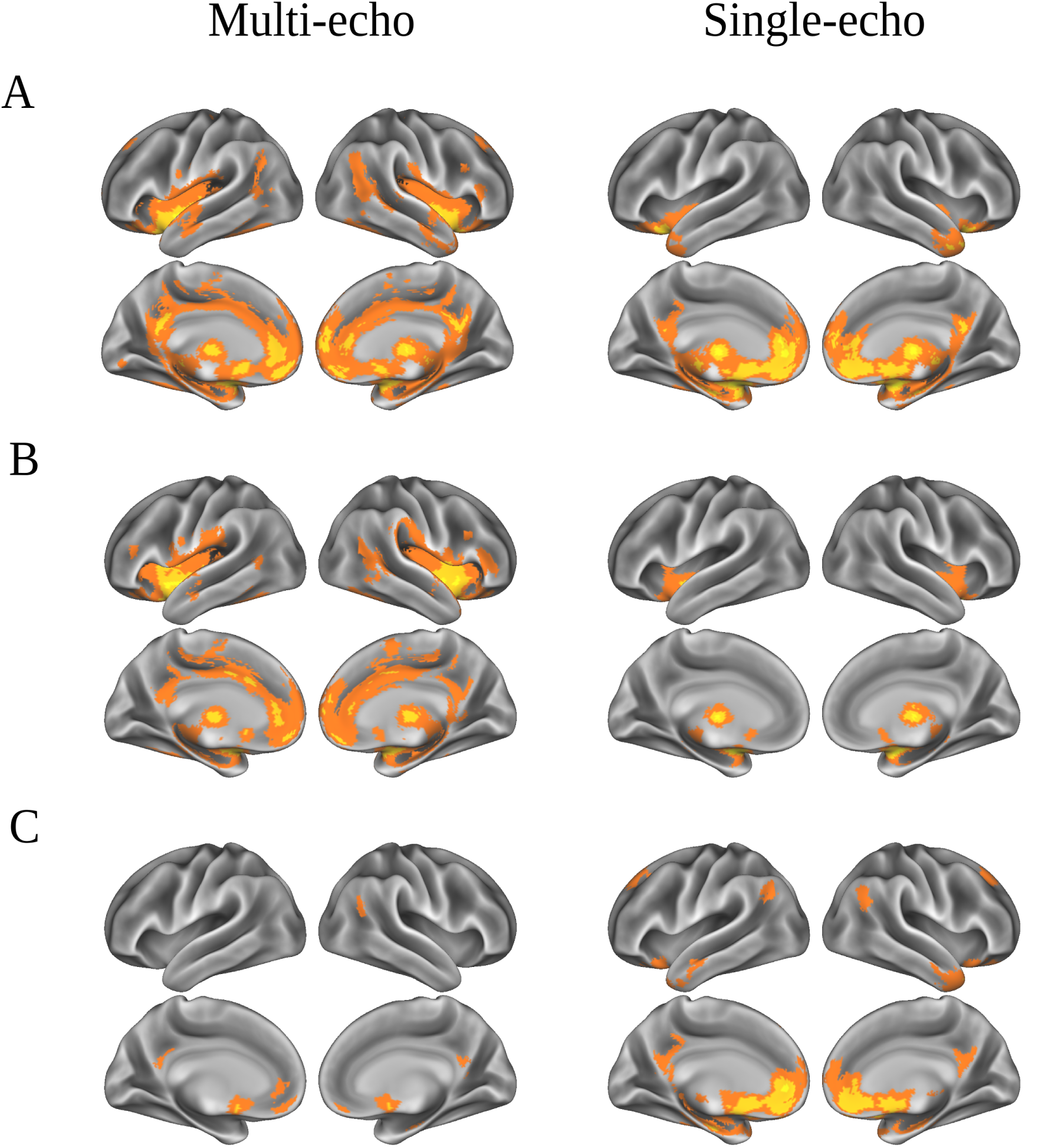
Inflated (i.e., surface) projections of multi-and single-echo functional connectivity results. **A**. Basal forebrain, **B**. NBM, and **C**. MS/DB ROI functional connectivity. Figure was generated using the Connectome Workbench software (Marcus et al., 2011).

#### 3.3.2 Distinct NBM and MS/DB connectivity

Examining functional connectivity of the Ch4 group of the BF (NBM) using ME-ICR revealed a similar profile to that of the entire BF ROI (Figure 4A, blue), with relationships to prefrontal cortex, the cingulum band, insulo-opercular cortex, fusiform gyrus, and subcortical features like the amygdala. Two additional clusters in the left inferior frontal gyrus and left postcentral gyrus were observed that had not been found in the whole BF analysis (Table 3).

**Table 3.** Nucleus basalis of Meynert (NBM) functional connectivity. Regions functionally connected with the NBM as assessed via ME-ICR (multi-echo) and time-series rsFC (single-echo). Coordinates are provided in MNI space. Sub-peaks were assessed within larger clusters based on a |*Z*| > 10 threshold and a minimum distance ≥ 10mm from other sub-peaks; locations for sub-peaks are italicized.

|  | *x* | *y* | *z* | *k* | *Z* |
| --- | --- | --- | --- | --- | --- |
| *Multi-echo results* |  |  |  |  |  |
| Right ventral striatum | 23 | 10 | -5 | 32704 | 12.69 |
| *Left putamen* | -23 | -8 | -3 |  | 12.57 |
| *Right caudate nucleus* | 13 | 16 | 3 |  | 11.63 |
| *Left caudate nucleus* | -11 | -12 | 5 |  | 11.39 |
| *Right putamen* | 33 | -4 | -5 |  | 11.09 |
| *Right thalamus* | 7 | -14 | 9 |  | 10.64 |
| *Right insula* | 41 | 10 | -5 |  | 10.60 |
| *Left thalamus* | -7 | -16 | 9 |  | 10.15 |
| *Left amygdala* | -23 | 0 | -15 |  | 10.07 |
| *Left insula* | -39 | 6 | -7 |  | 10.03 |
| Left anterior cingulate cortex | -3 | 42 | 11 | 7947 | 8.24 |
| Left middle temporal gyrus | -57 | -8 | -13 | 151 | 4.38 |
| Right middle frontal gyrus | 41 | 16 | 27 | 151 | 4.54 |
| Left middle temporal gyrus | -55 | -62 | 7 | 101 | 4.29 |
| Left middle frontal gyrus | -39 | 32 | 17 | 75 | 4.34 |
| *Single-echo results* |  |  |  |  |  |
| Right ventral striatum | 23 | 10 | -5 | 9080 | 12.63 |
| *Left putamen* | -23 | 8 | -5 |  | 12.42 |
| *Right amygdala* | 25 | -2 | -17 |  | 11.40 |
| *Left amygdala* | -21 | -4 | -15 |  | 11.27 |
| *Right caudate nucleus* | 15 | 4 | 15 |  | 10.57 |
| *Right thalamus* | 5 | -12 | 9 |  | 10.19 |
| *Left thalamus* | -3 | -14 | 7 |  | 10.11 |
| Right cerebellum (nodule) | 1 | -58 | -33 | 1888 | 7.41 |

Compared to ME-ICR, single-echo analyses showed more circumscribed connectivity for the NBM (Figure 4B, blue). Connections were noted with insulo-opercular cortex and subcortical targets, extending into the cerebellum (Table 3).

In contrast to the NBM, functional correlations from the Ch1-3 group of the BF (MS/DB) were more localized (Figure 4A, yellow). Multi-echo processed data revealed distinct connections between the MS/DB and the caudate (Table 4).

**Table 4.** Medial septum/diagonal band of Broca (MS/DB) functional connectivity. Regions functionally connected with the MS/DB as assessed via ME-ICR (multi-echo) and time-series rsFC (single-echo). Coordinates are provided in MNI space. Sub-peaks were assessed within larger clusters based on a |*Z*| > 10 threshold and a minimum distance ≥ 5mm from other sub-peaks; locations for sub-peaks are italicized.

| | $x$ | $y$ | $z$ | $k$ | $Z$ |
| --- | --- | --- | --- | --- | --- |
| *Multi-echo results* |  |  |  |  |  |
| Medial septal nucleus | -5 | 2 | -11 | 635 | 7.25 |
| Right medial frontal gyrus | 5 | 48 | -17 | 229 | 4.29 |
| Right superior temporal gyrus | 49 | -58 | 19 | 144 | 4.27 |
| Right parahippocampal gyrus | 27 | -16 | -19 | 121 | 4.18 |
| Left posterior cingulate cortex | -5 | -52 | 25 | 106 | 3.87 |
| Left anterior cingulate cortex | -5 | 46 | 1 | 85 | 4.04 |
| Right posterior cingulate cortex | 5 | -56 | 21 | 83 | 3.94 |
| *Single-echo results* |  |  |  |  |  |
| Medial septal nucleus | 11 | 12 | -11 | 10564 | 12.20 |
| *Left caudate nucleus* | -7 | 12 | -9 |  | 12.06 |
| *Left anterior cingulate cortex* | -5 | 46 | -1 |  | 10.38 |
| *Left hippocampus* | -25 | -14 | -19 |  | 10.37 |
| *Hypothalamus* | 7 | 2 | -11 |  | 10.21 |
| Left posterior cingulate | -7 | -52 | 17 | 784 | 4.99 |
| Left superior frontal gyrus | -19 | 40 | 43 | 263 | 5.64 |
| Left middle temporal gyrus | -61 | -10 | -17 | 232 | 4.42 |
| Right superior temporal gyrus | 51 | -60 | 27 | 189 | 4.66 |
| Right middle frontal gyrus | 21 | 32 | 47 | 156 | 4.73 |
| Left angular gyrus | -47 | -64 | 33 | 140 | 4.66 |

Single-echo processed data revealed broader connections from the MS/DB than were observed in multi-echo data (Figure 4B, yellow). Significant correlations included prefrontal cortex, medial temporal lobes, and posterior cingulate, with additional connections to the hippocampus and orbitofrontal cortex (Table 4).

## 4 Discussion

We assessed whole-brain, resting-state functional connectivity (rsFC) of the basal forebrain (BF) with multi-and single-echo fMRI data. The results demonstrate that the human BF supports broad functional connections with neocortical and subcortical regions, consistent with its prominent neuromodulatory role in nonhuman animals. Voxel-wise functional connectivity data from the BF were subjected to Ward clustering to examine functionally-defined BF nuclei, revealing almost perfect alignment between functional data and anatomical nuclei. Further rsFC analyses were conducted to examine the potential differential functional profiles of these anatomical nuclei. The nucleus basalis of Meynert (NBM) shows widespread functional correlations throughout lateral neocortex and subcortical regions, including the thalamus and amygdala, while connections with the medial septum/diagonal band of Broca (MS/DB) are more functionally connected to structures associated with the hippocampal formation.

### 4.1 Basal forebrain nuclei functional networks

The present results support a functional segregation of BF nuclei in the human brain. Clustering of human BF connections yielded outputs with high correspondence to anatomical delineations found in nonhuman animals. Moreover,the functional profiles were distinct for the NBM and MS/DB. Our results are in line with prior studies demonstrating a correspondence between anatomical connections and rsFC in humans and nonhuman primates (Genç et al., 2015; Hagmann et al., 2008; Honey et al., 2009; Margulies et al., 2009; Miranda-Dominguez et al., 2014; Mišić et al., 2016; Skudlarski et al., 2008).

Although the basal forebrain has a complex neurochemical profile (Lin et al., 2015), the functional connections from the MS/DB appeared to follow well-established cholinergic relationships, with observed correlations to the hippocampus, orbitofrontal cortex, and parahippocampal gyrus (Hedreen et al., 1984; Mesulam et al., 1983a,b). Although not directly assessed in the current study, the specificity of the functional profile revealed with these rs-fMRI data suggests that we may be observing relationships made through MS/DB cholinergic projections, rather than through the diffuse MS/DB GABAergic/glutamatergic network of projections (Mesulam et al., 1983b).

As hypothesized, we observed broad temporal correlations from the NBM to the amygdala and such neocortical regions as the insulo-opercular cortex, cingulum band, and prefrontal cortex. Homologous brain regions in rodent and nonhuman primate tracing studies have demonstrated similar strong cholinergic innervations from the NBM (Mesulam et al., 1983a, 1986, 1983b; Zaborszky et al., 2015a,b). Notable in their absence were functional connections with extrastriate cortical areas, which are also known to have anatomical connections with the NBM; however, this could be due to the nature of resting-state acquisition, for which no external stimuli were present, as this functional relationship has been observed in a previous human fMRI study during an experimental task with color stimuli (De Rosa et al., 2004).

### 4.2 Resting-state networks and the basal forebrain

Both the NBM and MS/DB showed distinct functional connectivity with regions typically associated with the default network (DN), including ventromedial prefrontal cortex (vmPFC), posterior cingulate cortex (PCC), angular gyrus, and the medial temporal lobes (Andrews-Hanna et al., 2014). While these regions are known to be innervated by BF projections, it appears BF targets do not fit neatly into the canonical human resting-state functional networks. Indeed, NBM rsFC analyses showed connectivity with regions that included the Default, Frontoparietal Control (FPCN; dorso-lateral prefrontal cortex), and Ventral Attention (VAN; insula) networks (Yeo et al., 2011). This is consistent with the BF as the source of a broad neuromodulatory network, playing a critical role across different facets of cognition and orchestrating information across these distinct resting-state networks.

### 4.3 Single-and multi-echo analysis

Despite previous reports of superior specificity of results and validity of statistical inferences with multi-echo fMRI data (Kundu et al., 2013, 2017), the current study suggests a more nuanced distinction between single-and multi-echo results. We compared multi-echo independent components analysis with a rigorous single-echo preprocessing pipeline (Power et al., 2012, 2014). Both the multi-and single-echo analysis pipelines detected discriminable functional networks from the BF nuclei (Figure 3); while multi-echo data seemed to provide greater detection power for the NBM functional network (Figure 4), the opposite appeared to be true for the MS/DB network (Figure 4).

It is worthwhile to note that the multi-echo preprocessing pipeline, which is entirely data-driven, retains many of the features of the original signal that are removed in the single-echo pipeline (i.e., by bandpass filtering, Gaussian smoothing) while still adequately attenuating noise (Figure 1). In this respect, multi-echo results may provide a truer snapshot of the underlying neural dynamics and connectivity profiles. We recommend future studies utilizing multi-echo data conduct joint analyses with single-and multi-echo processing pipelines to continue to provide a better picture of their unique benefits.

### 4.4 Methodological considerations

Our BOLD data, although likely reflecting underlying neural activity, are limited in lack of neurotransmitter specificity. Nonetheless, the observed connectivity patterns are consistent with known cholinergic projections of the BF (Gritti et al., 1994, 1997, 1998; Jourdain et al., 1989; Mesulam et al., 1983b; Meynert, 1872; Parent et al., 1988; Unal et al., 2015). Future studies could make direct assessments of cholinergic contributions by comparing network connectivity changes with systemic cholinergic pharmacology in humans.

### 4.5 Conclusions

Our current findings demonstrate differential functional correlation profiles of human BF nuclei. The NBM had diffuse connections with neocortical regions and the amygdala, while the MS/DB showed circumscribed relationships with the hippocampus and parahippocampal complex. These connections appear to support homologous anatomical relationships to the nonhuman rodent and primate literature (Mesulam et al., 1983a, 1986, 1983b). Given the importance of the BF as a key neuromodulatory hub, understanding these relationships is an important first step towards better characterizing the role of the BF in neural dynamics, its support of cognition, and its decline with neurodegeneration in aging.

## Acknowledgments

The authors wish to thank Elizabeth DuPre for helpful feedback on an original version of this manuscript. This work was supported by an Empire Innovation Program Grant [88021813] to E.D.R and an NIH NCRR grant [1S10RR025145] to the Cornell MRI Facility.

